# Population modeling with machine learning can enhance measures of mental health - Open-Data Replication

**DOI:** 10.1101/2022.04.04.487069

**Authors:** Ty Easley, Ruiqi Chen, Kayla Hannon, Rosie Dutt, Janine Bijsterbosch

## Abstract

Efforts to predict trait phenotypes based on functional MRI data from large cohorts have been hampered by low prediction accuracy and/or small effect sizes. Although these findings are highly replicable, the small effect sizes are somewhat surprising given the presumed brain basis of phenotypic traits such as neuroticism and fluid intelligence. We aim to replicate previous work and additionally test multiple data manipulations that may improve prediction accuracy by addressing data pollution challenges. Specifically, we added additional fMRI features, averaged the target phenotype across multiple measurements to obtain more accurate estimates of the underlying trait, balanced the target phenotype’s distribution through undersampling of majority scores, and identified data-driven subtypes to investigate the impact of between-participant heterogeneity. Our results replicated prior results from Dadi et a (2021) in a larger sample. Each data manipulation further led to small but consistent improvements in prediction accuracy, which were largely additive when combining multiple data manipulations. Combining data manipulations (i.e., extended fMRI features, averaged target phenotype, balanced target phenotype distribution) led to a three-fold increase in prediction accuracy for fluid intelligence compared to prior work. These findings highlight the benefit of several relatively easy and low-cost data manipulations, which may positively impact future work.

## 1. Introduction

In recent years, studies with sufficiently large participant samples (N>2000) have reported very low effect sizes for associations between neuroimaging measures and trait-level phenotypes (Dadi et al., 2021; Dutt et al., 2021; Marek et al., 2020). For example, Dadi et al used Random Forest Regression in N=10,000 participants from the UK Biobank (UKB) data to predict trait-level phenotypes of neuroticism and fluid intelligence, and reported a maximum R^2^ of 0.04 when using neuroimaging features (Dadi et al., 2021). Despite explaining a small amount of variance, results from these large participant samples are robust against sampling variability (Marek et al., 2022) and have good out-of-sample generalizability (Dadi et al., 2021; Dutt et al., 2021). Nevertheless, low brain-phenotype effect sizes are somewhat surprising given the presumed brain basis of phenotypic traits such as neuroticism and fluid intelligence. The goal of this paper is to replicate the findings from Dadi et al (Dadi et al., 2021) in a larger UKB sample and test multiple hypotheses regarding potential factors that may influence low brain-phenotype effect sizes. Specifically, we test whether addressing potential data pollution challenges (De Nadai et al., 2022) such as noisy phenotypes, noisy neuroimaging measures, skewed (imbalanced) phenotypic distributions, and population heterogeneity lead to improvements in R^2^. We focus on resting state functional MRI (rfMRI) neuroimaging measures to control the scope of this work, but relative improvements in R^2^ are expected to generalize to other modalities.

The first potential explanation for low brain-phenotype effect sizes is that the neuroimaging features used for prediction are noisy measures of brain function. Previous work reported UKB test-retest reliability results for neuroimaging measures ranging from 0.3 for rfMRI measures to 0.9 for structural measures (Dutt et al., 2021). The relatively lower test-retest reliability of rfMRI measures likely results from both dynamic state fluctuations and measurement error due to the relatively small number of timepoints (t=490). An interesting question is therefore whether residuals from the task fMRI scan (after removing task activation effects) can be combined with rfMRI data to reduce measurement error and improve prediction accuracy. This possibility is supported by previous work showing shared trait-level connectivity information between task residual and rfMRI data, although state-related connectivity differences were also observed (Fair et al., 2007). Furthermore, it may be possible to improve prediction accuracy by leveraging other types of features that can be extracted from rfMRI and task residual data (J. Bijsterbosch et al., 2020; J. D. Bijsterbosch et al., 2021). For example, network amplitudes have higher test-retest reliability than connectivity information (Dutt et al., 2021), and have been shown to capture individual differences in behavioral traits (J. Bijsterbosch et al., 2017; Miller et al., 2016). We test whether adding task residual connectivity features and/or including additional amplitude features will result in a higher R^2^.

The second potential explanation for low brain-phenotype effect sizes is that the target phenotype for prediction may be a noisy measure of the underlying trait. This potential explanation is supported by the fact that Dadi et al reported a substantially larger maximum R^2^ of 0.52 for age (compared to R^2^<0.04 neuroticism and fluid intelligence) (Dadi et al., 2021). Importantly, the phenotype of age is known without error (apart from potential database entry mistakes), whereas neuroticism and fluid intelligence scores are obtained from self-report questionnaires and test questions respectively, which are prone to intra-individual response instability, and potential social desirability bias in the case of neuroticism (McKelvie, 2004). These factors are reflected in previous estimates of test-retest reliability of neuroticism (0.85 in UKB (Dutt et al., 2021)) and fluid intelligence (0.65 in UKB (Lyall et al., 2016)). Response instability can be reduced by averaging over multiple available repeats of the same measure, thereby reducing noise and obtaining a more accurate estimate of the underlying trait. We test whether using phenotypes that have been averaged across all available UKB instances as the prediction target will result in a higher R^2^.

The third potential explanation for low brain-phenotype effect sizes is that the distribution of the target phenotype may be skewed, leading to an imbalanced regression problem (Yang et al., 18--24 Jul 2021). For example, neuroticism is strongly positively skewed such that the distribution peaks at zero (low N) with a long tail to the maximum score of 12 (high N; Fig. 1). Such underrepresentation of high neuroticism scores in population data will be propagated into the training sample, which likely results in inaccurate predictions of the underrepresented scores in the test sample, leading to a lower R^2^. A simple solution to this challenge is to flatten the distribution of neuroticism by undersampling the majority scores to ensure that all possible scores have equal representation (Dal Pozzolo et al., 2015). We test whether undersampling the UKB dataset to flatten the distribution of the target phenotype will result in a higher R^2^, despite the reduced overall sample size.

**Figure 1.**
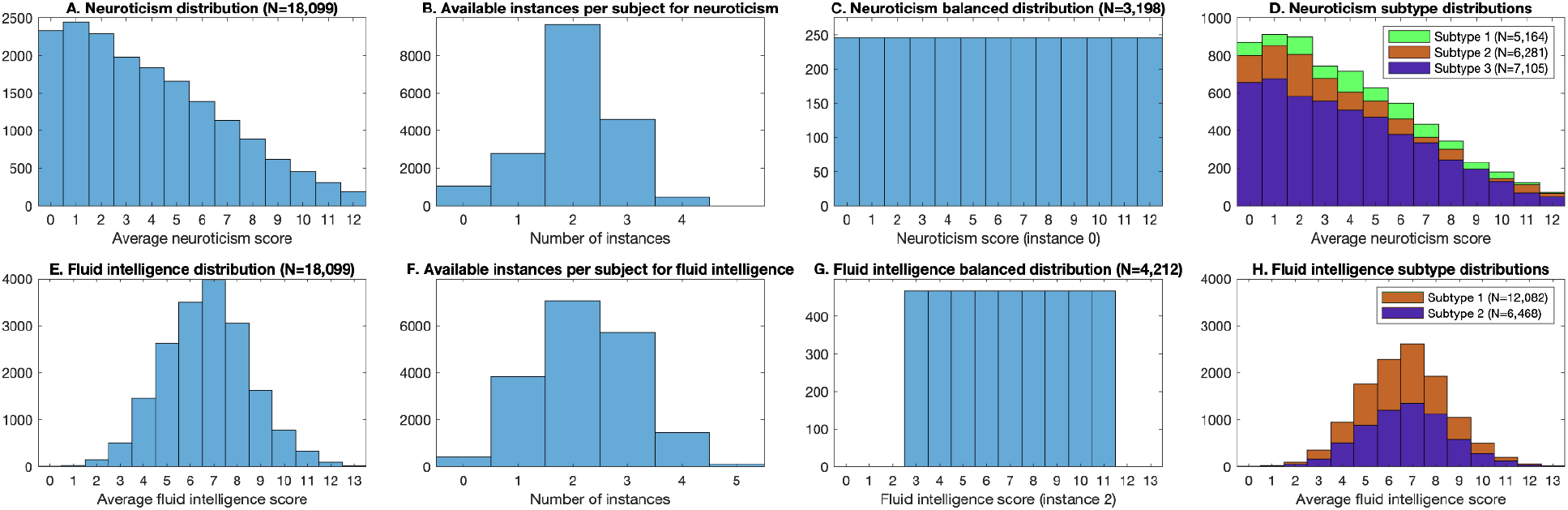
Target phenotype distributions. Top row (A-D) shows neuroticism and bottom row (E-H) shows fluid intelligence. The first column (A, E) displays the distribution of averaged phenotypes, the second column (B, F) displays the number of available instances per subject, the third column (C, G) displays the balanced phenotype distributions, and the fourth column (D, H) displays the distributions of data-driven subtypes.

The fourth potential explanation for low brain-phenotype effect sizes is potential between-participant biological heterogeneity. For example, two individuals with the same high score for neuroticism may not share the same brain-basis for their high neuroticism. Hence, different subgroups of participants may have divergent brain-phenotype associations, suggesting the presence of biological subtypes (also known as ‘biotypes’). If multiple biotypes are combined in the same analysis, the overall population effect size estimate may be diluted or canceled out (Ferrante et al., 2019). We adapt a previously developed data-driven pipeline to identify biotypes (Drysdale et al., 2017) based on canonical correlation analysis (Hotelling, 1936; Winkler et al., 2020) combined with rigorous cross-validation to test for biotype stability (Dinga et al., 2019). We test whether subsequently performing phenotype predictions separately within each biotype will result in a higher R^2^, despite the reduced sample size.

Taken together, the goal of this study is to better understand potential data pollution drivers of low effect sizes for brain-phenotype associations and to identify avenues for improvements to inform future research. Strengthening brain-phenotype associations is important for personalized medicine efforts and to maximize the impact of research into the brain basis of clinical, cognitive, and behavioral traits of interest (Paulus & Thompson, 2019, 2021). Our results reveal small but consistent increases in R^2^ from each of the tested avenues for improvements, which when combined resulted in a boost in R^2^ from 0.03 to 0.06 for fluid intelligence and from 0.00 to 0.03 for neuroticism.

## 2. Methods

### 2.1 Sample

Data from the UKB 20,000 neuroimaging release were used for this study. N=18,550 participants had complete resting state and task neuroimaging data, which is approximately double the participants used in Dadi et al (Dadi et al., 2021). The demographic characteristics were 51.6% female (5,572) and 48.3% male (5,403), and an age range at the time of scanning of 44–80 years (mean ± standard deviation: 62.5 ± 7.5 years). The sample was split in half to generate separate datasets for model training and model generalization. This research was performed under UK Biobank application number 47267.

### 2.2 Target phenotypes

Neuroticism was calculated for each instance using a summary score based on the sum across 12 binary (yes/no) questions (Table 1) developed by (Smith et al., 2013). This is identical to the field ID 20127 used in (Dadi et al., 2021), but enables calculation of Neuroticism scores for each of the available instances.

**Table 1.** UKB variables used as target phenotypes. Field IDs 1920-2030 were used to calculate neuroticisms. Each question is answered yes (1) or no (0) and the neuroticism score represents the sum across all responses. Field IDs 20016 and 20191 are identical measures of fluid intelligence.

| Phenotype | Field ID | Question/ description |
| --- | --- | --- |
| Neuroticism | 1920 | Does your mood often go up and down? |
|  | 1930 | Do you ever feel 'just miserable' for no reason? |
|  | 1940 | Are you an irritable person? |
|  | 1950 | Are your feelings easily hurt? |
|  | 1960 | Do you often feel 'fed-up'? |
|  | 1970 | Would you call yourself a nervous person? |
|  | 1980 | Are you a worrier? |
|  | 1990 | Would you call yourself tense or 'highly strung'? |
|  | 2000 | Do you worry too long after an embarrassing experience? |
|  | 2010 | Do you suffer from 'nerves'? |
|  | 2020 | Do you often feel lonely? |
|  | 2030 | Are you often troubled by feelings of guilt? |
| Fluid Intelligence | 20016 | Unweighted sum of the number of correct answers given to the 13 fluid intelligence questions completed on touch screen during assessment center visit |
|  | 20191 | Unweighted sum of the number of correct answers given to the 13 fluid intelligence questions completed in web-based online follow-up |

Within the UKB, the same fluid intelligence test was repeated at each assessment center visit and as part of a cognitive function online follow-up questionnaire. (Table 1). Both these variables provide an unweighted sum of the number of correct answers given to the same set of 13 fluid intelligence questions. Participants were given 2-minutes to answer the questions, and those who did not finish scored zero for each of the un-attempted questions.

Participants with missing data for individual questions resulting from answers such as ‘do not know’ or ‘prefer not to answer’ were removed for neuroticism and fluid intelligence. Specifically, neuroticism was only calculated for participants and instances with complete data on all 12 questions. For replications of original work by Dadi et al (Dadi et al., 2021), we used the target phenotypes of 20016 at instance 2 for fluid intelligence, and the sum neuroticism score at instance 0 (corresponding to 20127).

#### 2.2.1. Phenotype averaging

To test whether averaging the target phenotypes improved prediction accuracy, neuroticism and fluid intelligence scores were averaged across all instances available for each participant respectively. The range of available instances per participant was 0-4 for neuroticism and 0-5 for fluid intelligence (Figure 1).

#### 2.2.2. Phenotype flattened distribution

To test whether flattening the phenotype distributions improved prediction accuracy, we calculated the number of participants with the least common value (neuroticism = 12 for 246 participants, and fluid intelligence = 11 for 468 participants) and randomly selected the matching number of participants for each score. To achieve a sufficiently large sample size, fluid intelligence was truncated to range from 3 to 11 (inclusive) for the flat distribution tests (i.e., participants with fluid intelligence score 0, 1, 2, 12, 13, 14 were excluded). The total size of the flat distribution samples was N=3,198 for neuroticism and N=4,212 for fluid intelligence.

### 2.3 Resting state features

Preprocessed resting state data was downloaded from the UKB, as described previously (Alfaro-Almagro et al., 2018; Miller et al., 2016). The task residual data was not released with the UKB task data to limit the download size, so we repeated the fit of the task general linear model for each participant and saved the residuals. Previous work performed a group ICA on processed rfMRI data of N=5,000 UKB participants at a dimensionality of 100, out of which 55 components represented signal resting state networks (Miller et al., 2016). Dual regression was performed separately for rfMRI and task residual data after normalization to MNI space using all 100 canonical group components to obtain participant-specific resting state time series (Nickerson et al., 2017). From the 55 signal component dual regression time series, we estimated covariance matrices using Ledoit-Wolf shrinkage (Ledoit & Wolf, 2004) and used tangent-space embedding to transform the matrices into a Euclidean space (Dadi et al., 2019; Ng et al., 2014; Pennec et al., 2006; Pervaiz et al., 2020; Sabbagh et al., 2019; Varoquaux et al., 2010). We then vectorized the matrices’ lower triangles, obtaining 1,485 connectivity features.

#### 2.3.1 Combining rfMRI and task residual data

We separately calculated the feature vectors from rfMRI and task residual data and performed joint tangent-space embedding. To test whether combining rfMRI and task residual connectivity information improved prediction accuracy, we compared the effect of averaging the resulting feature vectors and of concatenating the vectors, doubling the feature space to 2,970 features.

#### 2.3.2 Adding amplitude features

To estimate the network amplitudes, we calculated the standard deviation of each signal network dual regression time series (J. Bijsterbosch et al., 2017), resulting in 55 additional features per scan per participant.

### 2.4 Predictive model

We used code released on GitHub by (Dadi et al., 2021) to implement random forest regression: https://github.com/KamalakerDadi/empirical_proxy_measures, which implements Random Forest Regressor in scikit-learn. Nested 5-fold cross-validation was used to tune the depth of the trees and the number of variables considered for splitting, and the tree depth was fixed to 250 trees. The full sample was split into random halves to generate a validation dataset for model construction and a held-out generalization dataset. Within the validation dataset, the Monte Carlo (or shuffle-split) resampling scheme was used to subdivide the validation set into 100 training (90%) and testing (10%) splits. In the generalization dataset, predictions were generated for each of the 100 models from each cross-validation split. Split validation performance was used to generate the violin plots.

### 2.5 Biotype definition

Data-driven clustering approaches to identify subtypes can be driven by phenotypic information (but the resulting subtypes might not differ in neuroimaging measures) or by rfMRI features (but the resulting subtypes might not be related to the phenotype of interest). To address this challenge, we adopted a method that enables brain-phenotype interactions to drive the definition of subtypes. To this end, Canonical Correlation Analysis (CCA, (Hotelling, 1936)) was used to relate the individual questions that make up the phenotypes of neuroticism and fluid intelligence to the rfMRI features (Drysdale et al., 2017), and the canonical scores were subsequently used to drive K-means clustering to identify biotype clusters. The pipeline included feature selection, CCA fitting and permutation testing, estimation of the optimal number of clusters using bootstrapping, and cluster assignment as explained in further detail below. The full pipeline was independently repeated across 100 cross-validations each using 90% of the data, and the Adjusted Rand Index (ARI) was calculated using shared participants for each pair of the 100 cross-validations to assess cluster stability (Dinga et al., 2019; Varol et al., 2017). The pipeline was finally performed on the full dataset using optimal solutions for feature selection and the number of clusters across 100 cross-validations to assign participants to each biotype. The predictive model was then separately applied in each biotype.

#### 2.5.1 Feature selection

Feature selection was performed prior to CCA keeping the top 25% of features to ensure a ratio of approximately 50 participants per feature, which is important to ensure robustness of the CCA (Helmer et al., 2021). To this end, separate unpaired t-tests were performed for each individual question making up the phenotype to compare participants who scored 0 (corresponding to ‘no’ in neuroticism and ‘incorrect’ in fluid intelligence) to participants who scored 1 (corresponding to ‘yes’ in neuroticism and ‘correct’ in fluid intelligence). The absolute t-statistics were summed across all questions within the phenotype and the 25% of features with the highest combined t-statistic were entered into the CCA (i.e., 371 features out of 1,485). Feature selection was repeated within each cross-validation fold. The 25% features that we most commonly selected across the 100 cross-validations were used in the final run on the full dataset.

#### 2.5.2 CCA and permutation testing

CCA was performed between the selected rfMRI features and all separate questions within the phenotype using permCCA (Winkler et al., 2020). Permutation testing was performed (2,000 permutations) to identify significant canonical covariates using family wise error (FWE) correction to control for multiple comparisons across the canonical covariates. For all canonical covariates with a FWE-corrected p-value below 0.05, the canonical scores were used for subsequent clustering. Each canonical score has two values per participant related to rfMRI features and phenotype questions respectively (which are correlated because this correlation represents the canonical correlation). Both canonical scores were used for clustering. CCA fitting and permutation testing was repeated within each cross-validation fold.

#### 2.5.3 Estimation of optimal number of clusters

Bootstrapping with replacement was performed 1,000 times to test cluster solutions with k=2-10. For each bootstrap, the k with the highest silhouette score was recorded. The mode across all 1,000 bootstrap was subsequently implemented as the optimal number of clusters. Estimation of the optimal number of clusters was repeated within each cross-validation fold. The optimal number of clusters was that occurring most commonly across the 100 cross-validations and was set in the final run on the full dataset. K-means clustering with the optimal number of clusters was performed to assign each participant to a cluster. K-means clustering was performed separately in each cross-validation fold and in the full dataset. To assess the stability of the clusters, the Adjusted Rand Index (ARI; implementation from McComb, https://github.com/cmccomb/rand_index; (Vinh et al., 2010)) was computed from the cluster assignment for each of the 100 cross-validation folds.

## 3. Results

### 3.1 Replication of previous results

Our replication of both neuroticism and fluid intelligence using a larger UKB sample resulted in slight increases in R^2^ (0.01 vs 0.00 for neuroticism; 0.03 vs 0.02 for fluid intelligence; Fig. 2) for both phenotypes compared with previous work (Dadi et al., 2021). These results are consistent with the gradual monotonic increase with sample size shown in supplementary figure 1 of Dadi et al (Dadi et al., 2021).

**Figure 2.**
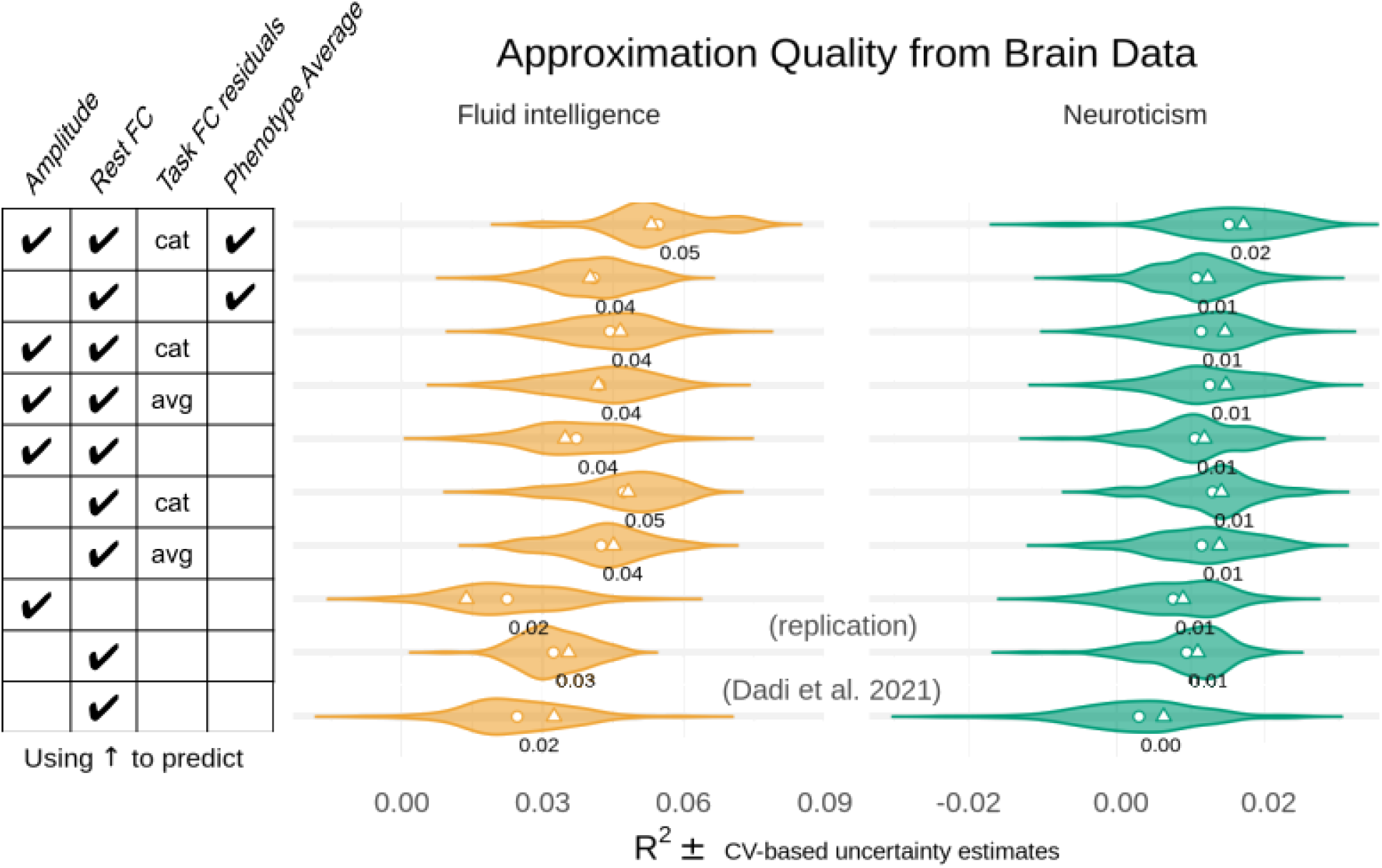
Including additional resting state features leads to improvements in prediction accuracy. We report the R^2^ metric to facilitate comparisons across prediction targets. The cross-validation (CV) distribution (100 splits) on the validation dataset is depicted by violin plots. Circles depict the average performance on the validation data across CV-splits, and triangles depict the performance of the average prediction (CV-bagging) on held-out generalization datasets. For convenience, the mean performance on the validation set is written for each violin plot. In the table on the left ‘cat’ indicates that resting state and task connectivity features were concatenated and ‘avg’ indicates that resting state and task connectivity features were averaged.

### 3.2 Effect of including additional fMRI features

The inclusion of additional fMRI features yielded modest increases in R^2^ for fluid intelligence and small increases in R^2^ for neuroticism. For fluid intelligence, each additional fMRI feature prompted a small increase in R^2^, from 0.03 to 0.04, except in the case of concatenation (Fig. 2). Concatenating tangent-space projected residual task features provided the largest increase in R^2^ of any single-feature intervention for fluid intelligence (0.05 vs. 0.03; Fig. 2). No change in average prediction accuracy for neuroticism was observed until all interventions were employed. When fMRI features were combined and used to predict average neuroticism, R^2^ doubled from 0.01 to 0.02 (Fig. 2). Fluid intelligence also saw its largest increase in prediction accuracy in the presence of all interventions: R^2^ increased from 0.03 to 0.05 (Fig. 2). Using only signal amplitudes as features yielded prediction accuracies similar to those on full connectivity data in smaller sample sizes (0.02 vs. 0.02 in fluid intelligence; 0.01 vs. 0.00) in prior work (Dadi et al., 2021). However, rest-amplitude and amplitude-only predictions showed a larger decrease in generalization performance relative to validation performance when compared to other methods (Fig. 2).

### 3.4 Effect of phenotype averaging

The use of average phenotype values as prediction targets yielded small increases in prediction accuracy for fluid intelligence and neuroticism. Targeting average fluid intelligence prompted an improvement in prediction accuracy for both rest-only (from 0.03 to 0.04; Fig. 2) and all-feature (from 0.04 to 0.05; Fig. 2) fMRI feature sets. By contrast, average neuroticism only yielded an improvement in R^2^ in the all-feature case (0.01 to 0.02).

### 3.5 Effect of flattened (balanced) phenotype distribution

Selecting subjects to generate a flat distribution of phenotype prediction targets yielded slight increases in R^2^ for fluid intelligence (0.04 vs. 0.03, rest; 0.06 vs 0.05, all features), but not for neuroticism (0.00 vs 0.02, all features; 0.01 vs. 0.00, rest) when compared with predictions on the same features (Figure 3).

**Figure 3.**
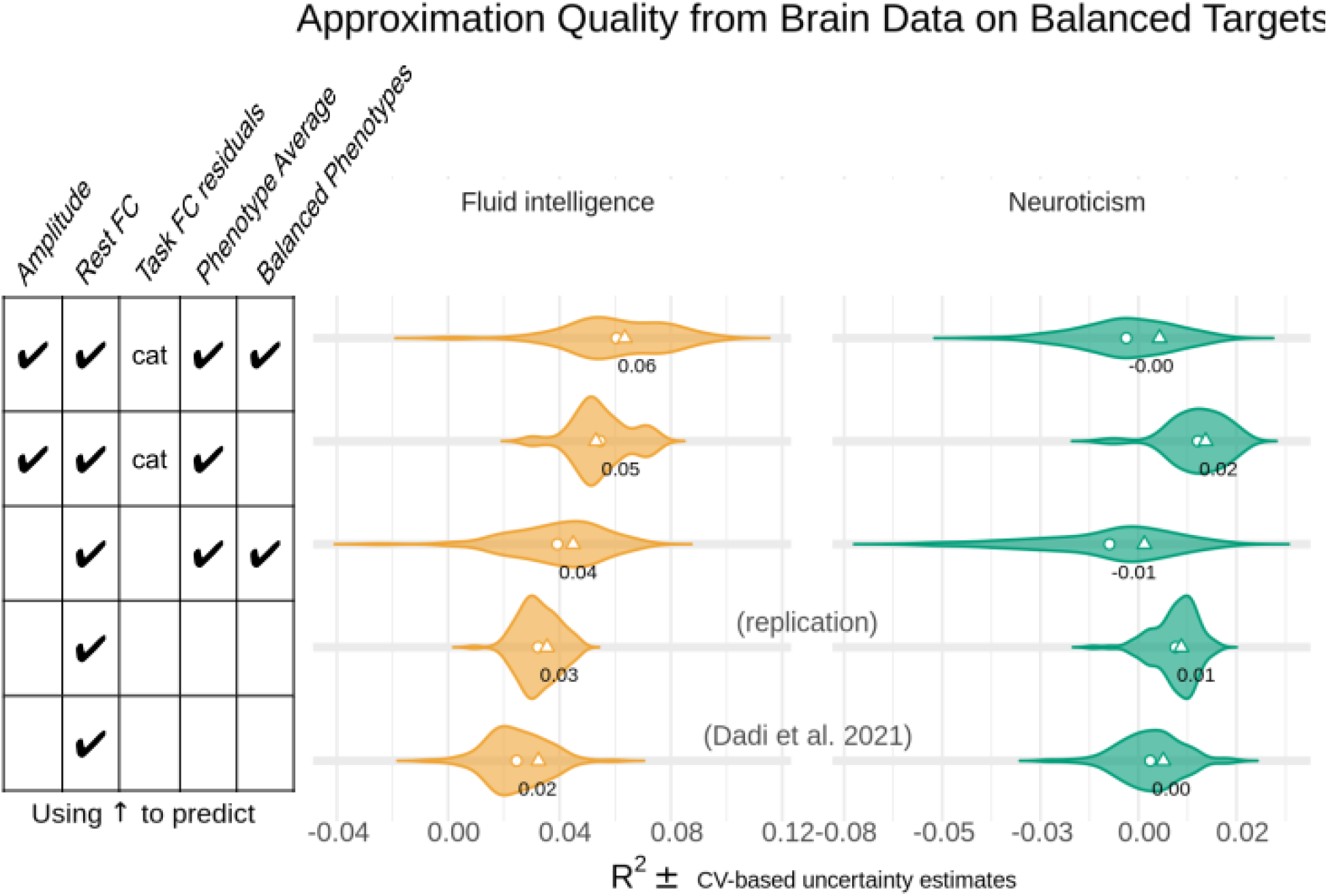
Averaging the phenotypes and balancing the sample led to additional improvements in prediction accuracy for fluid intelligence, over and above improvements achieved by including additional resting state measures. We report the R^2^ metric to facilitate comparisons across prediction targets. The cross-validation (CV) distribution (100 splits) on the validation dataset is depicted by violin plots. Circles depict the average performance on the validation data across CV-splits, and triangles depict the performance of the average prediction (CV-bagging) on held-out generalization datasets. For convenience, the mean performance on the validation set is written for each violin plot. In the table on the left ‘cat’ indicates that resting state and task connectivity features were concatenated.

### 3.6 Effect of separate predictions in biotypes

Distributions of canonical correlations and their *p*-values are taken over the 100 cross-validation valids; family-wise error corrected *p*-values are computed from 2000 permutations within each cross-validation fold. The 95% confidence intervals for the means and *p*-values of 0.05-significant canonical correlations (after family-wise error correction) are given in Table 2. Confidence intervals clear the significance threshold (i.e., are bounded above by 0.05 after FWE correction) in four canonical components in fluid intelligence and two in neuroticism. Across the 1000 bootstrap iterations for k-means clustering, the mode cluster number with the highest silhouette score was two for fluid intelligence and three for neuroticism. The Adjusted Rand Index (ARI) across cross-validation folds indicated moderate stability for both the two fluid intelligence clusters (ARI=0.76), and the three neuroticism clusters (ARI=0.72). Sample sizes for the resulting subtypes are reported in Fig. 1D and H.

**Table 2.** 95% confidence intervals for mean canonical correlations and their corresponding p-values after family-wise error correction. Four components of the fluid intelligence CCA were significant, and two components of the neuroticism CCA were significant.

|  | Quantity | Component 1 | Component 2 | Component 3 | Component 4 |
| --- | --- | --- | --- | --- | --- |
| Fluid intelligence | correlation | (0.214, 0.224) | (0.181, 0.193) | (0.173, 0.184) | (0.165, 0.170) |
|  | p-value | 0.0005 | 0.0005 | (0.0003, 0.0007) | (0, 0.004) |
| Neuroticism | correlation | (0.229, 0.242) | (0.200, 0.216) | (0.193, 0.205) | (0.188, 0.197) |
|  | p-value | 0.0005 | 0.0005 | (0, 0.2) | (0.5, 1) |

Repeating the random forest regression separately within each subtype did not result in improved prediction accuracy in either fluid intelligence or neuroticism. Prediction accuracy suffered in comparison to prediction from the full dataset (Fig. 4), and was comparable to accuracies observed in prior work on a smaller sample (Dadi et al., 2021).

**Figure 4.**
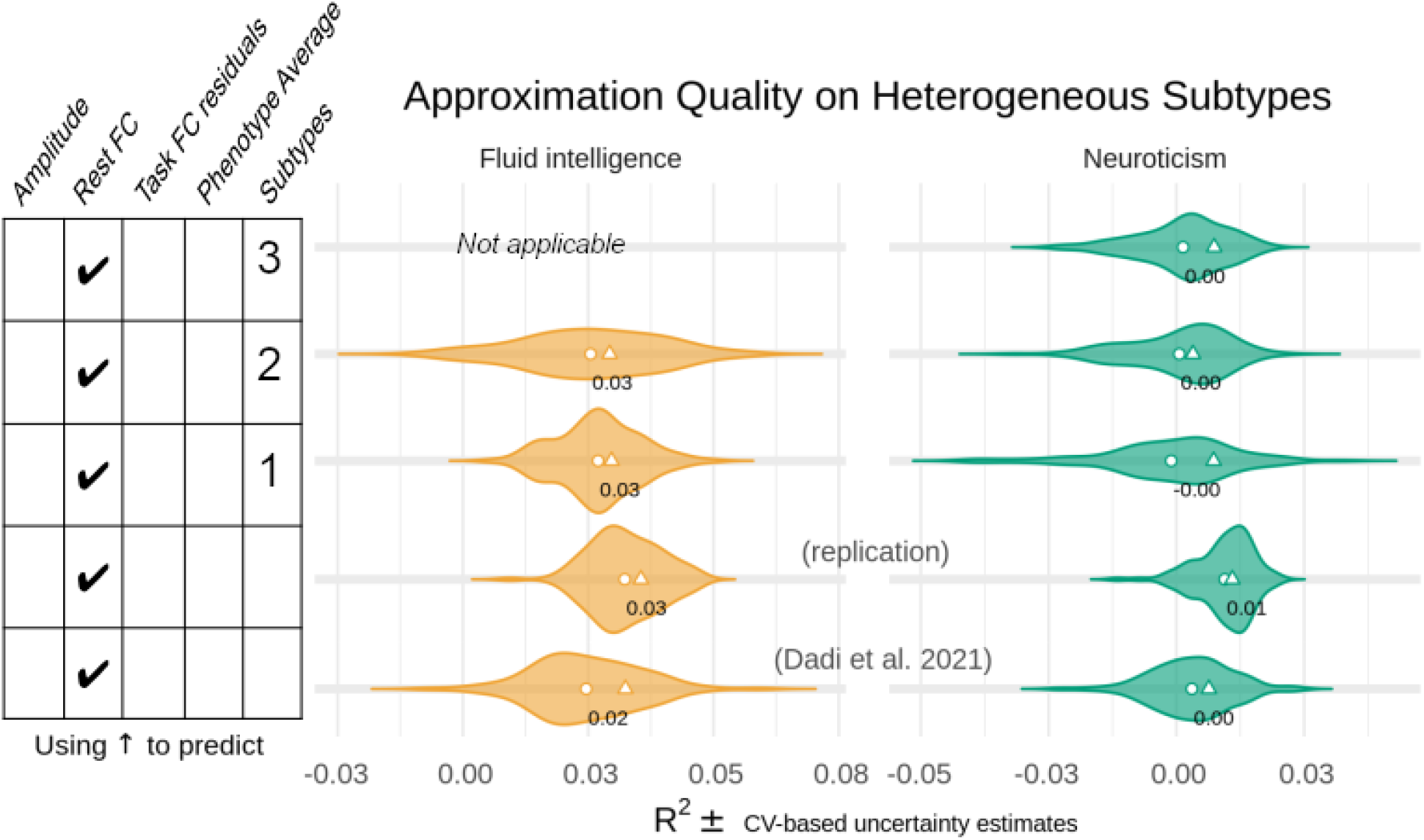
Identifying homogeneous subgroups did not result in major improvements in prediction accuracy. We report the R^2^ metric to facilitate comparisons across prediction targets. The cross-validation (CV) distribution (100 splits) on the validation dataset is depicted by violin plots. Circles depict the average performance on the validation data across CV-splits, and triangles depict the performance of the average prediction (CV-bagging) on held-out generalization datasets. For convenience, the mean performance on the validation set is written for each violin plot.

## 4. Discussion

In this paper we replicated and extended previous efforts (Dadi et al., 2021) to predict trait phenotypes of neuroticism and fluid intelligence based on rfMRI neuroimaging features. Our primary goal was to test different manipulations of the data to address data pollution challenges (De Nadai et al., 2022). Our replication findings using a larger sample achieved higher R^2^ without any further manipulations, pointing to the benefit of larger sample sizes even beyond N=10,000. Beyond this boost from sample size, our results revealed that each manipulation led to small but consistent increases in R^2^, which were largely additive when multiple manipulations were combined (Figures 2, 3, 4). The manipulations to address data pollution largely fall into three categories, namely: input fMRI feature data additions, target prediction phenotype averaging, and input participant changes. In the remainder of the discussion, we summarize the results and provide recommendations for each of these categories.

Firstly, we tested different manipulations of the input fMRI feature data by combining resting state and task residual datasets, and incorporating additional amplitude rfMRI features. Our findings showed small but consistent increases in R^2^ (Fig. 2). Of note, the amount of combined functional MRI data per person in the UKB (5 minutes rfMRI + 5 minutes tfMRI; (Miller et al., 2016)) is lower than other datasets such as the Human Connectome Project (1 hour rfMRI + 1 hour tfMRI; (Glasser et al., 2016)), and substantially lower than densely sampled datasets such as the Midnight Scan Club (5 hours rfMRI + 6 hours tfMRI; (Gordon et al., 2017)), so it is possible that further gains may be achievable with more data. Although, the UKB test-retest reliability estimates for rfMRI measures (Dutt et al., 2021) are in line with other datasets (Noble et al., 2019), suggesting that further gains may be limited. Although the addition of task fMRI features led to improvements, this manipulation requires substantially more time and computational resources than the other manipulations discussed below. Future work will need to carefully consider the trade-off between maximizing prediction accuracy and investment of time and other resources.

Secondly, we tested whether averaging the target phenotype for prediction across multiple measurements to obtain a less noisy estimate of the underlying trait improved R^2^. The results showed improvements, which were additive when combined with other manipulations (Fig. 2). Indeed, the largest improvement for neuroticism was obtained when combining extended fMRI features and averaging the target phenotype, which increased R^2^ from 0.0 in Dadi et al to 0.2 for Neuroticism. If multiple measurements are readily available (such as in the UKB), we recommend leveraging the data and using averaged phenotypes as an easy and low-cost manipulation that leads to consistent improvements in R^2^.

Thirdly, we tested two options to alter the participant sample by balancing the distribution of the target phenotype through undersampling the majority scores and by splitting the sample into separate biological subtypes. Notably, each of these approaches substantially reduced the sample size from 18,000 to samples ranging from 3,000 to 12,000 participants. Despite the reduced sample sizes, the balanced sample manipulation resulted in consistent improvements in R^2^ (Figure 3), highlighting the importance of a balanced sample over and above pure sample size. The largest improvement for fluid intelligence was obtained when combining extended fMRI features, averaging the target phenotype, and using a balanced sample, which tripled R^2^ from 0.2 in Dadi et al to 0.6 for fluid intelligence. Splitting the sample up into homogeneous subtypes did not result in major improvements in R^2^ (Fig. 4), which may be driven by smaller sample sizes and may point to residual heterogeneity from other sources. For example, alternative subtyping approaches that group participants based on the pattern of responses across individual questions may be beneficial.

In conclusion, we tested various data manipulations to address data pollution in an attempt to improve the prediction accuracy of neuroticism and fluid intelligence traits based on large-scale neuroimaging data from the UK Biobank. Each data manipulation led to a small but consistent increase in R^2^, and combining all data manipulations achieved a three-fold increase in R^2^ for fluid intelligence compared to prior work (Dadi et al., 2021). We recommend that future studies may implement phenotype averaging and target distribution balancing as relatively low-cost manipulations that lead to small but consistent improvements in prediction accuracy.

## 5. Code and Data Availability

All analysis code for this article will be made available shortly at: https://github.com/cmccomb/rand_indexhttps://github.com/PersonomicsLab/. UK Biobank data (Miller et al., 2016; Sudlow et al., 2015) are available following an access application process, for more information please see: https://www.ukbiobank.ac.uk/enable-your-research/apply-for-access.

## 6. Acknowledgements

We are grateful to UK Biobank and the UK Biobank participants for making the resource data possible, and to the data processing team at Oxford University for producing the shared processed data. This research was performed under UK Biobank application number 47267. This research was supported by the NIH (1 R34 NS118618-01) and the McDonnell Center for Systems Neuroscience.

